# Water motion as a transformation mechanism of algal communities structure in Lake Baikal

**DOI:** 10.1101/313650

**Authors:** Lyubov S. Kravtsova, Igor B. Mizandrontse, Svetlana S. Vorobyova, Lyudmila A. Izhboldina, Elena V. Mincheva, Tatyana G. Potyomkina, Tatyana I. Triboy, Igor V. Khanaev, Dmitry Yu. Sherbakovv, Andrey P. Fedotov

## Abstract

The diversity of algal communities of phytoplankton and meio-and macrophytes was investigated in Lake Baikal. Fragments of *Spirogyra* thallomes were recorded in the phytoplankton community of Southern Baikal, which had never been recorded before in its composition. It was also established that the structure of benthic algal communities changed in comparison with that in 2000 due to intense development of filamentous algae, particularly *Spirogyra*. Its lowest biomass was recorded in the surf zone and wave breaking, whereas the highest biomass was registered in the area of weakened effect of waves on the bottom. The cover percent of the bottom with filamentous algae in different areas of the coastal zone varied from 0 to 100%. Hydraulic characteristics of *Spirogyra* were the same as those of planktonic diatoms. The circulation currents and wave effect on the bottom favoured transfer and distribution of *Spirogyra* from the location of its intense development into the coastal area of Lake Baikal.

## Introduction

Alongside the effects of global warming and anthropogenic impact, algal blooms have been recorded in the coastal areas of seas and lakes (Nozaki *et al.* 2003; Smith *et al*. 2006; Hiraoka *et al.* 2011), which cause deterioration of not only water quality but also social environments in recreational areas. A similar phenomenon has been shown to exist in the ecosystem of Lake Baikal, a UNESCO World Heritage site. At present, large clumps of filamentous algae thrown to the shore have been found on the Baikal beaches in some areas of the lake (Timoshkin *et al*. 2016). Fragments of benthic filamentous algae *Spirogyra* have been found in the phytoplankton community (Kobanova *et al*. 2016; Bondarenko and Logacheva 2016), which have not been found in this community before (Kozhov 1931; Popovskaya 1977). It is necessary to determine the cause(s) of the emergence of *Spirogyra* cells in the phytoplankton.

For the past decade, changes have been recorded not only in the plankton structure (Izmest’eva *et al.* 2016) but also in the benthic communities of Lake Baikal (Kravtsova *et al*. 2014; Timoshkin *et al*. 2016). In the areas of the coastal zone confined mainly to settlements, we observe overgrowing of the bottom with filamentous algae among which there are members of the genus *Spirogyra*, something that is atypical of algal communities of Lake Baikal. Earlier, singular *Spirogyra* filaments have been recorded only in the well-heated bays and shallow areas (sors) of Lake Baikal and its tributaries (Kozhova and Izhboldina 1994; Izhboldina 2007). We hypothesise that the distribution of the members of the genus *Spirogyra* in the coastal zone of the lake is attributed to circulation currents of Baikal waters, which transfer these algae from places where they develop in great numbers. It is known that the hydrodynamic regime together with such environmental factors as temperature, light and chemical composition of water affect the biota structure of sea and freshwater environments (Peters *et al*. 2006; Wolcott 2007; Wang *et al*. 2012; Liu *et al*. 2015). Moreover, the water motion directly or indirectly influences hydrobionts. Specifically, characteristics of water motion, including velocity of currents, rough water, dynamic pressure and turbulence, directly affect mobility and transfer of aquatic organisms (Luchar *et al*. 2010; Durham *et al*. 2013; Cross *et al*. 2014). Wind waves and ripples affecting higher aquatic plants and benthic communities may also function as limiting and optimising factors in their growth (Raspopov *et al.* 1990). An indirect effect of water motion on the diversity and spatial distribution of hydrobionts occurs due to the changes of sedimentation dynamics, transport of particulates and detritus in the coastal area of water bodies (Snelgrove *et al*. 1988; Airoldi 1998).

It is interesting to know how hydrodynamics affect the flora of Lake Baikal, a unique freshwater environment with a depth of over 1,630 m combining the features of sea and lake ecosystems. The objective characteristic of water motion in Lake Baikal has been obtained from long-term in-situ measurements, instrumental investigation and mathematical models. General patterns of formation of currents, wave activity, surging, turbulence and upwelling were determined previously (Pomytkin 1960; Ainbund 1973; Afanasyev and Verbolov 1977; Fialkov 1983; Zhdanov *et al.* 2009; Shimaraev *et al.* 2012). There are few works on the indirect effect of hydrodynamics on the biota of Lake Baikal, which are devoted only to the survey of the effect of water masses on plankton (Likhoshway *et al.* 1996; Jewson *et al*. 2010) and wave activity on distribution of benthic organisms in the coastal area of the lake (Karabanov and Kulishenko 1990).

The aim of this study is to assess the role of *Spirogyra* in the structure of current algal communities of Lake Baikal and contribution of hydrodynamic processes in its dissemination in the coastal zone of the lake.

## Material and methods

### Field studies

We studied the algal flora of Lake Baikal in August 2016 on board the research vessel “Titov”. We analysed phytoplankton of Southern Baikal to study the possible transfer of *Spirogyra* fragments (Fig. 1*a*). Phytoplankton samples (1.5 L of water) were collected at six stations (I–VI) with a water sampler at depths of 0, 5 and 15 m. Stations I–V were located at a distance of approximately 50–100 m off the shore. Station VI was 7 km away from the shore in the direction from Cape Listvennichny to the settlement of Tankhoy. Additionally, we collected samples from depths of 25 m and 50 m. All quantitative phytoplankton samples (22) were fixed in Utermel solution.

**Fig. 1.**
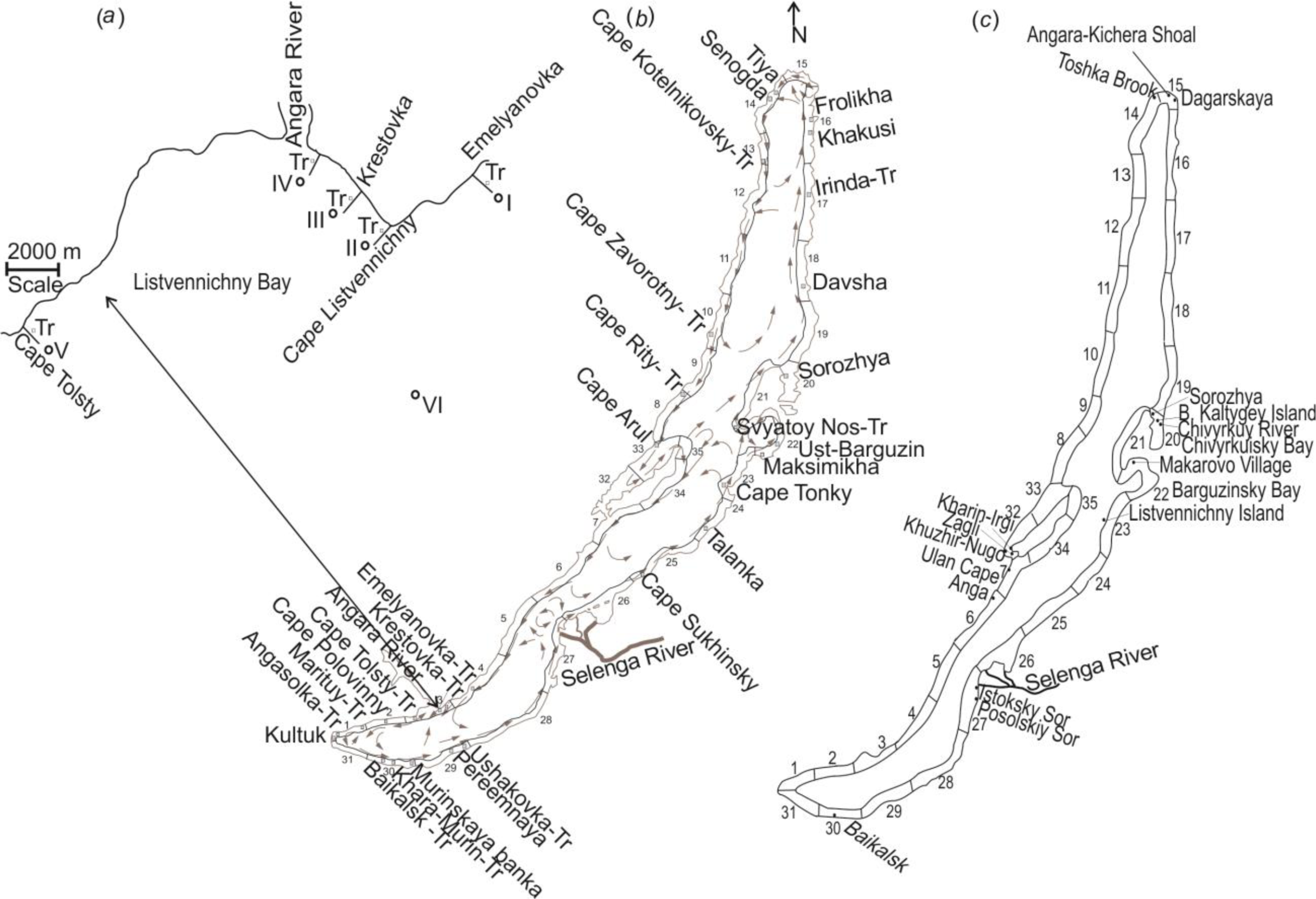
(*a,b*) Map-scheme of algal sampling in different areas of Lake Baikal and (*с*) areas of findings (before 2000) of singular *Spirogyra* filaments. (*a*) I-VI – stations of phytoplankton sampling in Listvennichny Bay; (*b*) – geographic names on the scheme corresponding to points of quantitative algal samples at stations; Tr – transects at which scuba divers measured bottom cover percent with filamentous algae (%) and simultaneously collected samples; 1-35 – stations differing in wind-wave characteristics and bottom geomorphology (according to Fialkov 1983); arrows show circulation of water masses (according to Afanasyev and Verbolov 1977); (*с*) – points show places of *Spirogyra* findings (archive data of L.Izhboldina).

To estimate the recent diversity of phytobenthos, scuba divers collected 117 quantitative samples of meio- and macrophytes from depths of 0–10 m in Southern, Central and Northern Baikal at 29 stations located at 11 sites differing in wind-wave characteristics and bottom geomorphology (Fig. 1*b*). Moreover, to assess the bottom cover percent with filamentous algae, the scuba divers mapped meio- and macrophytes at 15 transects (Tr) using frames (area of 1 m^2^) divided into 100 equal quadrats.

The scuba divers also collected 18 quantitative samples of meio- and macrophytes from two transects directed perpendicular to the shoreline to characterise the structure of benthic algal communities in Southern Baikal. One transect was located in the background region (outside the impact zone of the settlement of Listvyanka) 5 km to the north of Cape Listvennichny opposite Emelyanovka Valley. Another transect was located in the impact zone in Listvyanka opposite Krestovka Valley (Fig. 1*a*). The scuba divers collected three samples in each vegetation belt at depths of 0–10 m using a frame with an area of 0.16 m^2^. The scuba divers put stones covered with algae into sacks of strong fabric and lifted them onboard the vessel. The stones were then put into a large cuvette with water. The algae were cut with a scalpel from the surface or brushed off. The water was poured into the sieve of a mill-gauze No. 23. The algae were put into flasks and fixed in 4% formalin.

#### Laboratory analysis

Phytoplankton samples were settled in a 15–20 mL volume for about 14 days. Algae were counted in 0.1 mL. Individual volumes of cells were taken into account to determine algal biomass (mg m^−3^) (Makarova and Pichkily 1970). Picoplankton and cysts of chrysophytes were not considered in total phytoplankton biomass.

To assess the transport in currents of both *Spirogyra* and diatoms, we calculated the sinking velocity during settling in a laminar flow from Stokes formula:

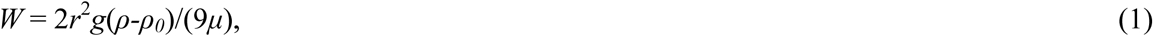

where *W* is the sinking velocity of a spherical particle in the water, cm s^−1^; *r* is its radius, cm; *g* is the gravity acceleration (normal, 980.655), cm s^−2^; *ρ* is the particle density, g cm^−3^; *ρ_0_* is the water density (1 g cm^−3^); *μ* is the water dynamic viscosity, g cm^−1^ s^−1^.

To reduce non-spherical algal cells to a conditionally spherical shape, we calculated the equivalent radius according to cell volume:

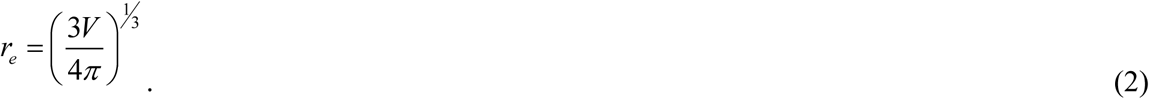

The geometric form coefficient *ζ* was used to estimate non-spherical particles:

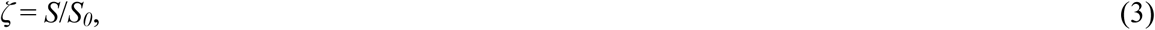

where *S* is the particle surface area; *S*_0_ is the sphere area of the same effective radius.

Dynamic coefficient of the particle shape *Г* (*ζ*) in the linear area of environmental resistance was calculated from Velikanov *et al.* (2013):

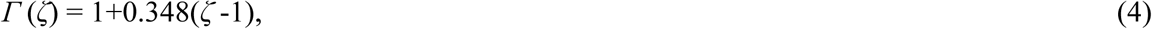

where *ζ* is the geometric form coefficient.

The density of living *Spirogyra* was determined in the following way. First, we estimated the true specific weight of dry mass (density of dry substance) of *Spirogyra* filaments. We dried them on filter paper and then subjected to a solid tablet to pressure in a compression mold. The density of the dry substance was calculated from the volume of this tablet and its weight. The humidity was estimated from the difference between the weight of *Spirogyra* filaments dried on filter paper (until a wet spot disappeared) and the weight of these filaments (quantity 20 g) dried at 103 °C.

The temperature of the cell content was set as equal to the water temperature for estimating the density of living *Spirogyra*. The water density at this temperature was determined from the table data. The density of living *Spirogyra* was calculated from the following formula:

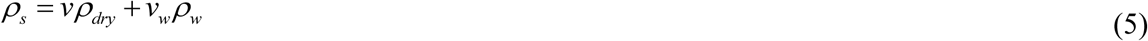

where *v* and *v_w_* are the volumes of the dry substance and water, respectively; *ρ_dry_* is the density of the dry substance; *ρ_w_* is the water density at the given temperature.

Meio- and macrophytes were sorted according to taxon level under an MBC 10 microscope at 2×8 magnification. Species were identified from the temporary algal preparations under an Amplival microscope at 12×10 and 12×40 magnifications. Cell sizes (diameter, width and length in μm) were measured with an ocular-micrometer.

Before weighing meio- and macrophytes on a torsion balance VT-500 with an accuracy of 0.1, we dried them on filter paper until a wet spot disappeared. The data obtained were converted to 1 m^2^ of the bottom (g m^−2^).

The diversity of phytoplankton as well as of phytobenthos was characterised using Shannon’s species diversity index (Odum 1971):

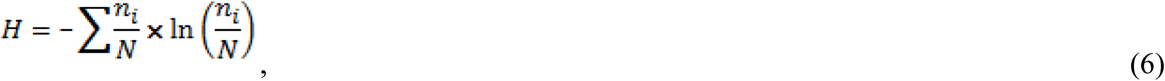

where *n_i_* is the biomass of *i*–species; *N* is the total biomass of species in a certain habitat.

Algal communities (of both phytoplankton and phytobenthos) were identified from the modified density index (Brotskaya and Zenkevich 1939):

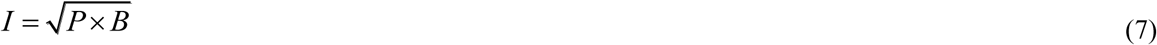

where *P* is the frequency of occurrence (the ratio of a number of samples in which a species has been found to total the number of samples, %); *B* is the percentage of a species in the total biomass, %. Species with maximal density index (*I*) were considered dominant, with *I* > 10% being subdominant and species with *I* < 10% being considered minor.

We compared the structure of algal communities with that of previous years, referring to the data by S. Vorobyova on phytoplankton in August 1992 (Southern Baikal – 35 quantitative samples) and L. Izhboldina on phytobenthos for August in 1966–1988 (Southern, Central and Northern Baikal – 37 quantitative samples out of 298).

The dependence of the lake bottom overgrowing with filamentous algae on the hydrodynamic environment was estimated by principal component analysis (PCA) using the following parameters as variables: x_1_ – bottom cover percent with filamentous algae, %; x_2_ – composition of bottom sediments; x_3_ – depth, m; x_4_ – width of the coastal zone, m; x_5_ – wave height (*h*, m); x_6_ – wave length (*λ*, m); x_7_ – periodicity of wave activity (*τ*, s); x_8_ – slope ratio of the coastal zone; x_9_ – bottom current velocity (*U*_max_ and *U*, m∙s^−1^); x_10_ – shear velocity (*V_sh_*, m∙s^−1^) of sediment movement (0.5 mm in diameter); x_11_ – coefficient of sediment mobility (*K_m_*); x_12_–x_18_ – hydrodynamic pressure (*P*, g m^−2^) on vertical surface at certain depths (0.5 m; 1.5 m; 2 m; 3 m; 4 m; 5 m). In our calculations, we used the highest values of wave activity (λ, τ) at the prevailing wind velocity (5–10 m∙s^−1^) during the August navigation (Galazy 1993).

During wave activity, the water moves around circular orbits. The sizes of these circulations decrease towards the shore (with the decrease in depth), and their orbits acquire the shape of flat ellipses. Moreover, bottom velocities of water motion increase. According to the interaction between the water flow and the bottom, we distinguish three zones during wave activity: I – offshore zone of undeformed waves; II – transformation of wave (deformation zone and breaker zone); and III – surf zone (Fig. 2*a*).

**Fig. 2.**
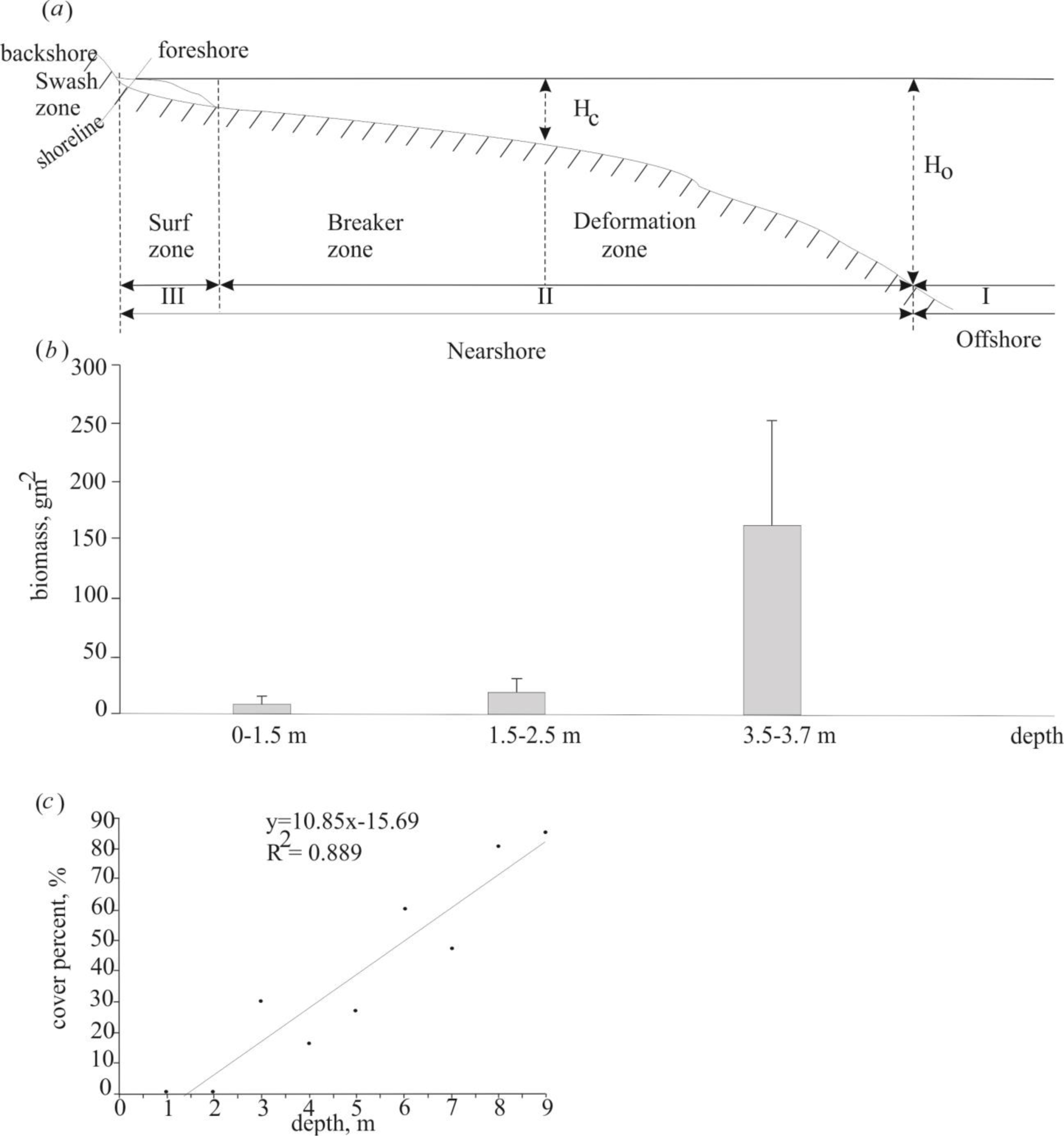
(*a*) Zones of wave effect on the bottom, (*b*) biomass of Spirogyra, and (*c*) correlation between cover percent with filamentous algae and depth in the coastal zone of Lake Baikal (Krestovka Valley, August 2016). *H*_c_ - a depth with wave effect on the bottom, and *H*_0_ - a depth without wave effect on the bottom.

The maximal velocity of the water motion (in the zone of undeformed waves) was calculated from the following equation (Petrov 1985):

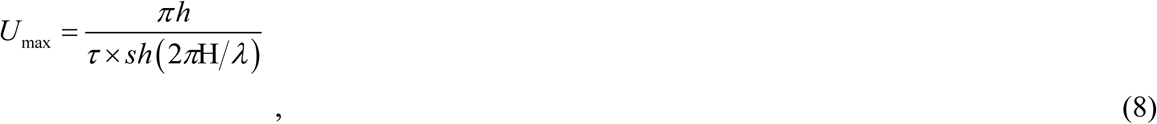

where *h* is the wave height, m; *τ* is the wave period, m; *H* is the depth, m; *sh* is the hyperbolic sine; λ is the wave length, m.

Current velocities were calculated from the following formula (Petrov 1985) in case of a significant effect of the bottom on the orbital wave component (in the zone of wave breaking):

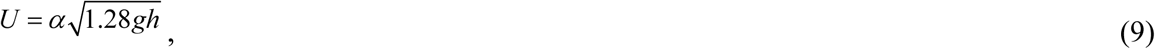

where *h* is the wave height, m; *g* is the gravity acceleration, m∙s^−2^; *α* is values varying from 0.7 to 1.8 with the depth decrease; in the surf zone it reaches 2 and then reduces to 0 at the end point of the splash.

The shear velocity (the initial velocity of sediment movement) was estimated from the following formula (Longinov 1963):

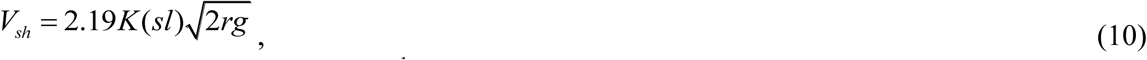

where *V_sh_* is the shear velocity, m∙s^−1^; *K(sl)* is the non-dimensional coefficient depending on the bottom slope (*sl*); *r* is the particle radius; *g* = 9.80665 m∙s^−2^:

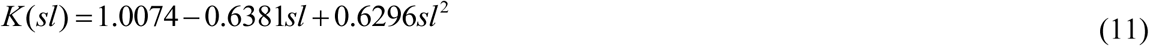

The mobility coefficient of bottom sediments was calculated from the ratio of the maximal horizontal component of the orbital bottom velocity and the initial velocity of sediment movement (Karabanov and Kulishenko 1990):

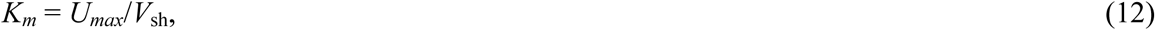

Water flow pressure on the vertical surface during wave activity was estimated from the following formula (Longinov 1963):

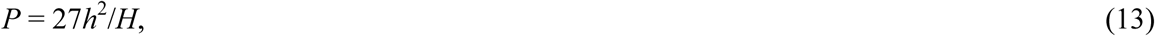

where *P* is the hydraulic pressure, g m^−2^; *h* is the wave height, m; *H* is the depth, m; numerical coefficient, g m^−2^ m^−1^.

## Results

During this study, the water temperature was 15–17 °C at a depth of up to 15 m in the coastal zone and in the adjacent areas of the open pelagic zone. In 2016, 50 taxa (at a lower level than genus) of planktonic algae were registered in the flora, of which 14 were diatom taxa, 3 were dinophyte taxa, 2 were cryptophyte taxa, 7 were chrysophyte taxa, 8 were blue-green taxa, 14 were green taxa and 2 were flagellate taxa. The diversity from the Shannon index varied from 2.6 to 2.8 in the coastal zone at the depths of 0 m, 5 m and 15 m. Algal communities represented by 38–44 taxa, including *Spirogyra* sp. with dominance of *Asterionella formosa* Hass., were found at these depths (Fig. 3*a*). Previously, in Southern Baikal in summer (1992), the Shannon index varied from 1.8 to 2.4 at different depths. At this time, the community comprised dominant species of *Rhodomonas pusilla* (Bachm.) Javorn. and *Gymnodinium coeruleum* Ant.; members of the genus *Spirogyra* were not recorded (Fig. 3*b*).

**Fig. 3.**
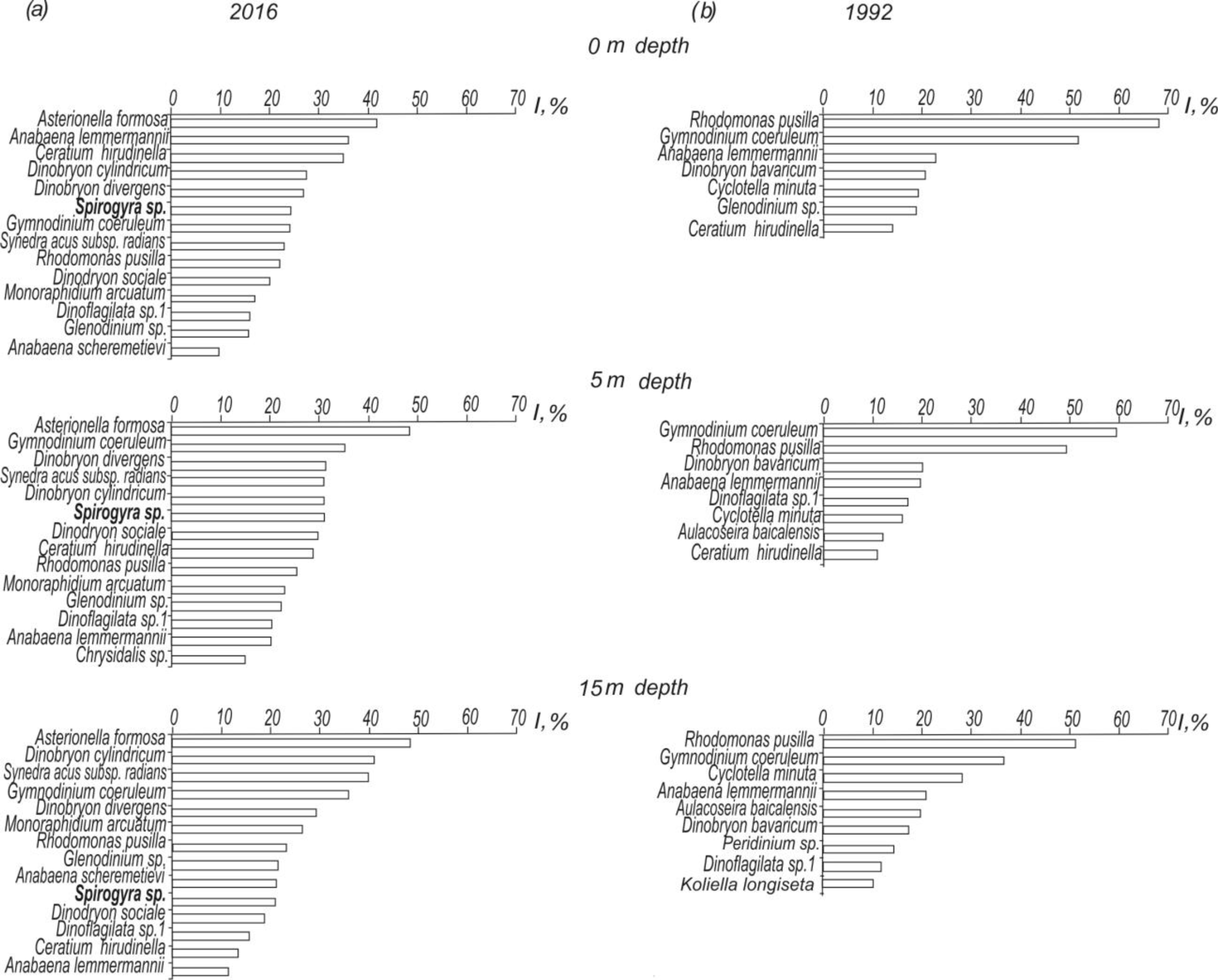
Structure of phytoplankton community in Southern Baikal during different years: (*a*) – in 2016; (*b*) – in 1992. Along X-axis – density index; along Y-axis – species ranking in the order of decrease of density.

At the depths of 25 m and 50 m in the open pelagic area, *Spirogyra* was also recorded in the phytoplankton community in 2016. *Fragilaria radians* Kütz., *Dinobryon cylindricum* Imhof., *R. pusilla* and *G. coeruleum* dominated. In 1992, the community was dominated by the latter two species distributed at these depths; *Spirogyra* was not recorded.

It should be noted that thallome fragments of *Spirogyra* consisting mainly of one to three cells were found in phytoplankton at all studied depths whose biomass varied from 8.5 to 35.2 mg m^−3^ at 0 m, 5 m, 15 m, 25 m and 50 m (the abundance was 650 to 5,440 cells L^−1^).

Geometric and hydraulic characteristics of both singular cells and thallome fragments of *Spirogyra* (Table 1) were assessed to ascertain possibilities of their transfer with coastal currents. The same characteristics were presented for comparison of diatoms: namely *A. formosa* as a dominant of the phytoplankton community and *Aulacoseira baicalensis* (Meyer) Simonsen as a typical member of the Baikal algal community. It was noted that the dynamic coefficient in the linear area of resistance for a *Spirogyra* cell and thallome fragments of several cells is comparable with that of diatoms. The density of dry *Spirogyra* calculated from the volume of the tablet and its weight was 1.36 g cm^−^ 3, and humidity of the living alga was 90%.

**Table 1.**
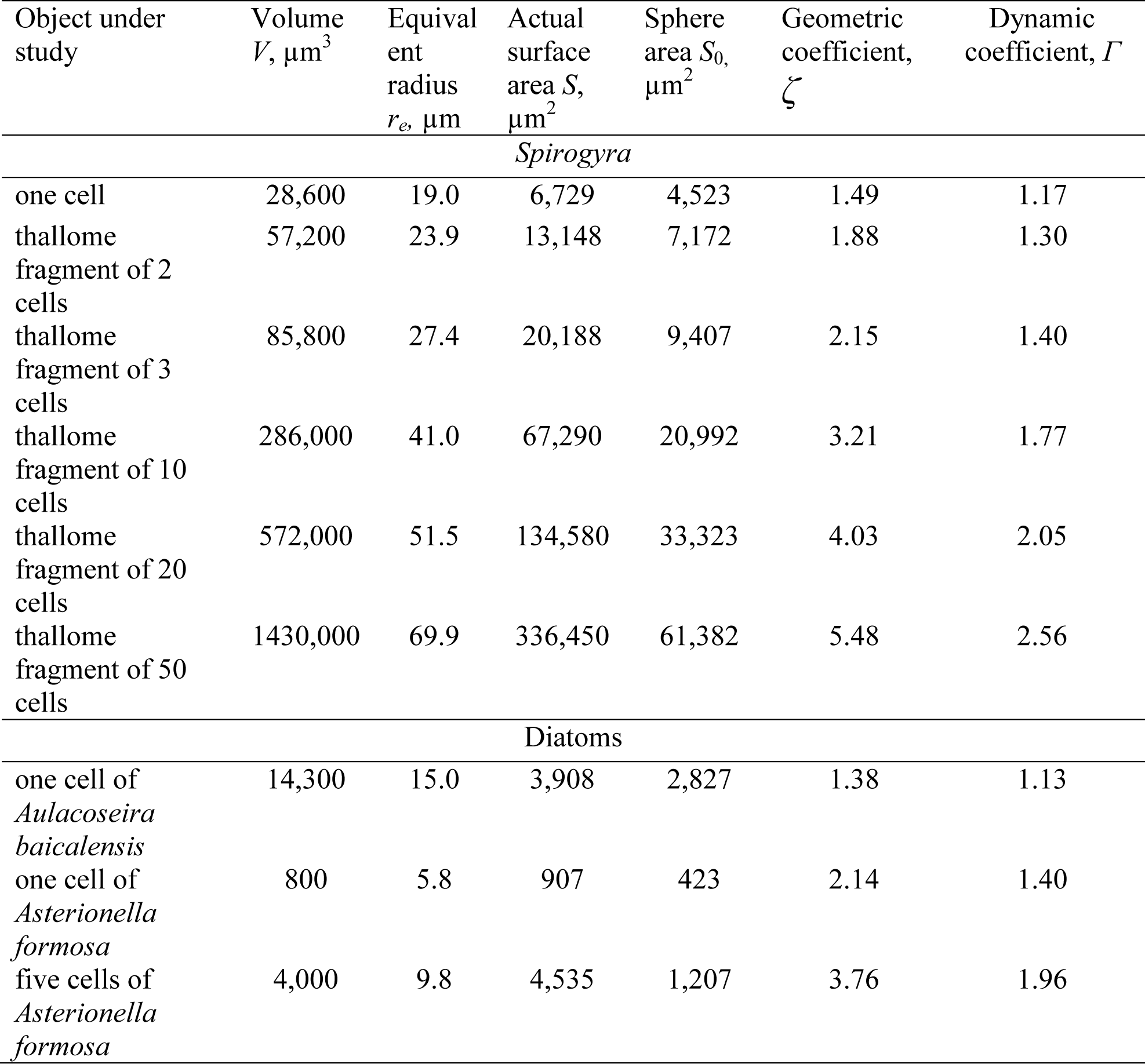
Geometric and hydraulic characteristics of *Spirogyra* and planktonic diatoms.

The density *ρ_s_* of the living *Spirogyra* was 1.036 g cm^−3^. The sinking velocity at 10 °C of one *Spirogyra* cell with the volume of 28,600 μm^3^ and equivalent radius of 19 μm was 22×10^−3^ mm∙s^−1^. The density of a thin-walled diatom *A. baicalensis* calculated by us from the data provided by Jewson *et al*. (2010) was 1.27 g cm^−3^. Therefore, its cell, with a volume of 14,300 μm^3^ and equivalent radius of 15 μm, sinks at a velocity of 90×10^−3^ mm∙s^−1^. Another diatom, *A. formosa,* sinks at 15×10^−3^ mm∙s^−1^ having a cell volume of 800 μm^3^ and equivalent radius of 5.76 μm. It is likely that this diatom is of the same density as that of *A. baicalensis*. As *A. formosa* is able to form star colonies consisting of several cells, e.g. five, its sinking velocity in this case is 44×10^−3^ mm∙s^−1^. The density of living skeletonless algae at this temperature is close to water density varying within small values. Even in such large cells as *Spirogyra,* the sinking velocity is lower than in a diatom of *A. baicalensis* with a silicon exoskeleton.

In 2016, the benthic flora in the studied regions of Lake Baikal was more diverse in comparison with that of the period before 2000. Its composition consisted of 65 taxa, 56 of them being meio- and macrophytes (Table 2), 8 higher aquatic plants (*Batrachium* sp., *Elodea canadensis* Michx., *Fontinalis antipyretica* Hedw.*, Myriophyllum* sp., *M. spicatum* L.*, Potamogeton crispus* L. and *P. perfoliatus* L.) and a lichen *Collema ramenskii* Elenk.

**Table 2.**
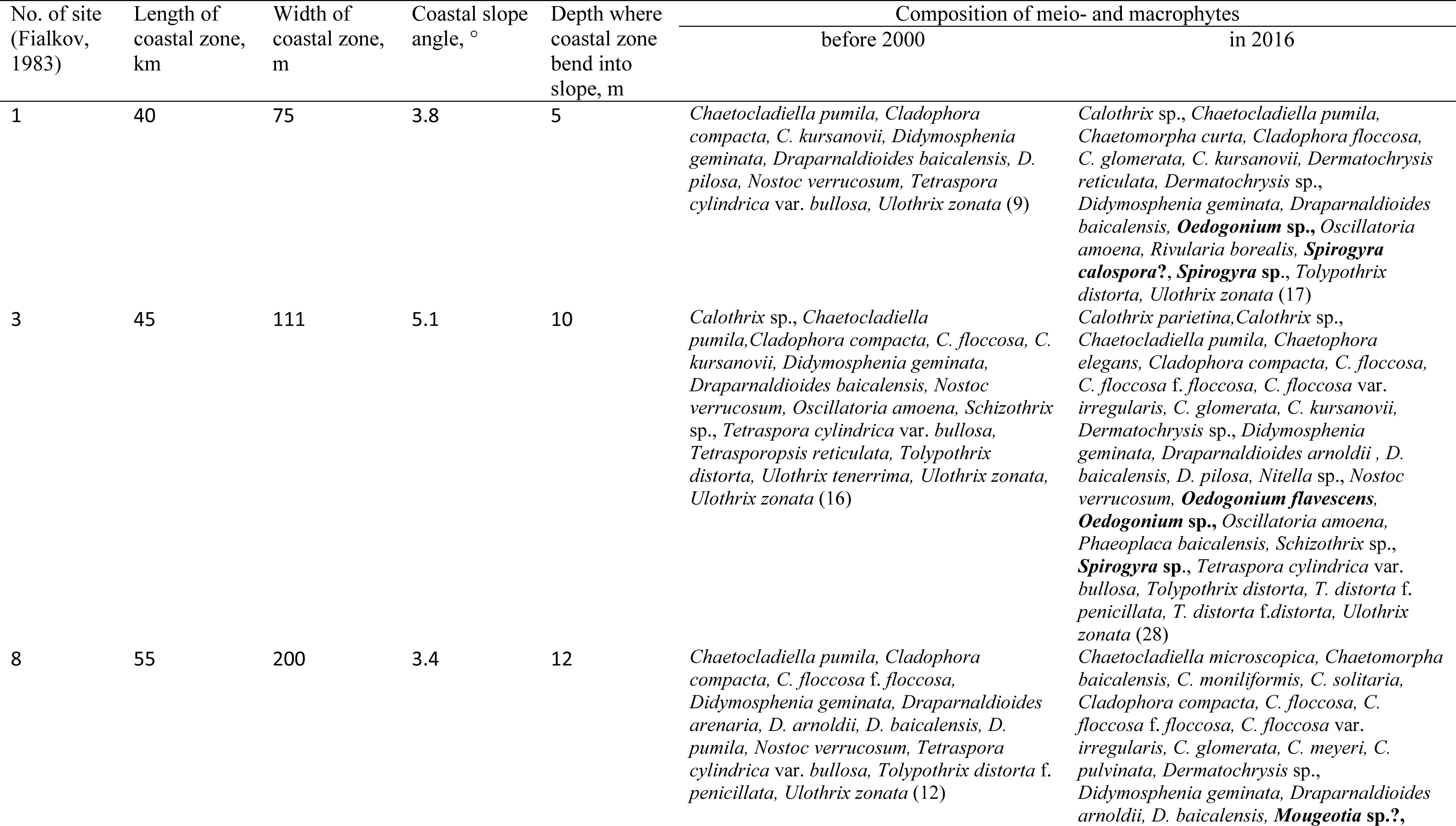
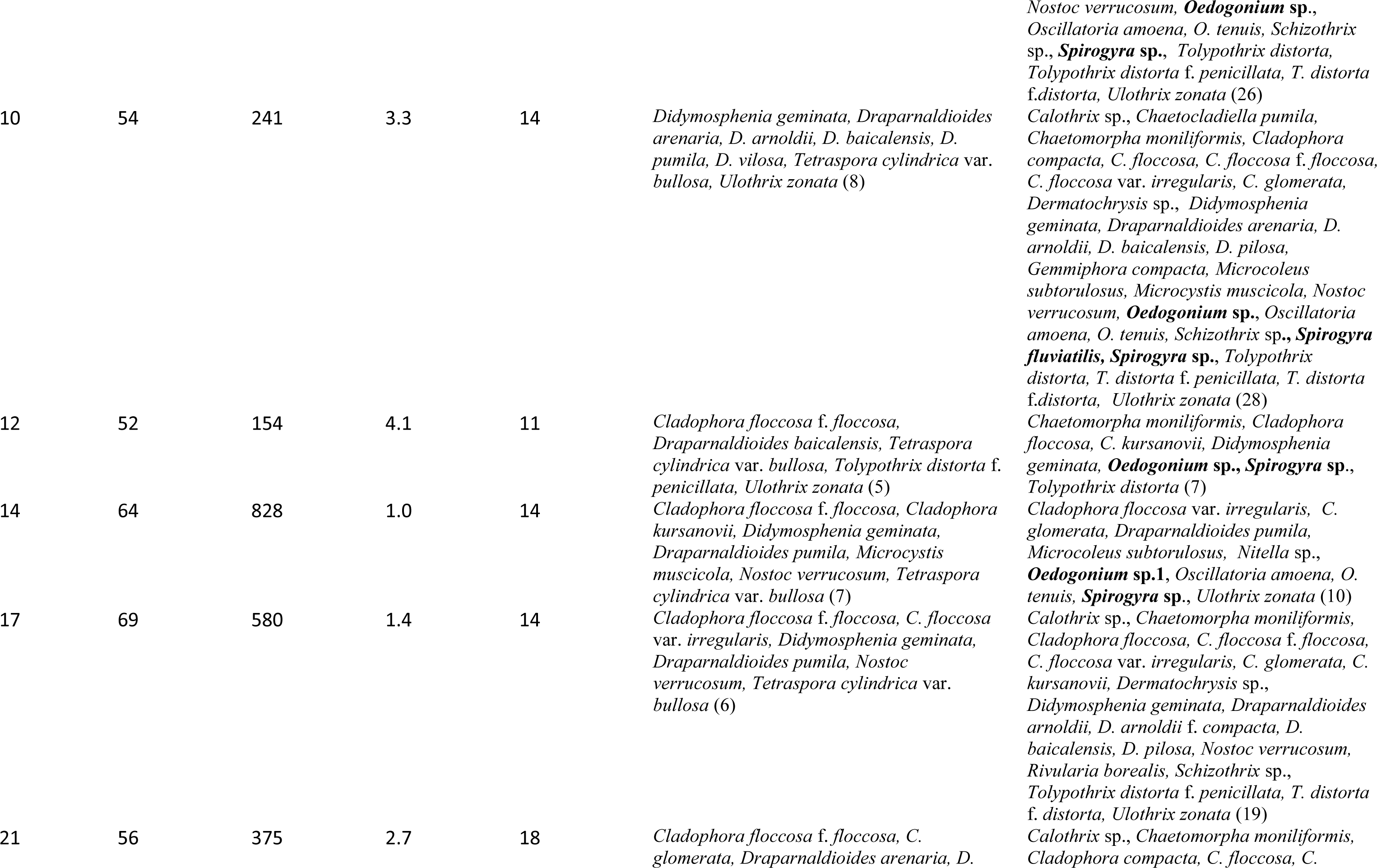
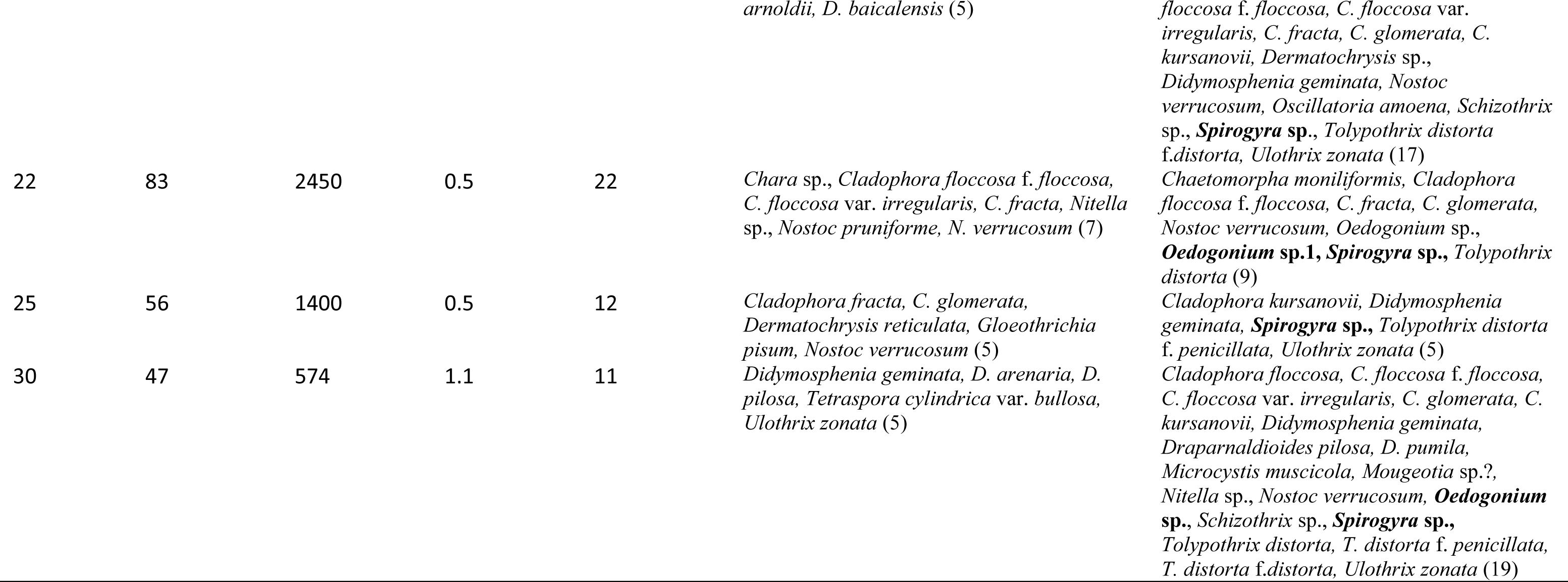
Composition of meio- and macrophytes in the coastal zone of Lake Baikal. Taxa uncharacteristic of the coastal zone in bold; total taxon number in brackets

Among meio- and macrophytes, we detected the filamentous algae *Mougeotia* sp., *Oedogonium flavascens*?, *Oedogonium* sp., *Spirogyra calospora*?, *Spirogyra* sp. and *Ulothrix zonata* (Web. et Mohr.) Kütz. Of special interest were algae of the genus *Spirogyra* whose habitat in the previous century was confined only to the coastal-sor zone (Fig. 1*c*), whereas at present the habitat has widened significantly (Fig. 1*b*). Before 2000, *Spirogyra* inhabited only certain areas of the lake in the form of singular filaments (Fig. 1*c*). Moreover, it was recorded in the grab samples collected at a depth of 40–80 m (Dagarskaya Bay, Angara-Kichera Shoal and Cape Ukhan). In 2016, the occurrence of *Spirogyra* was 75% along the open coastal areas in Lake Baikal. Besides *Spirogyra*, filamentous algae of the genus *Oedogonium* were also widespread in the lake with an occurrence of 40% (Table 2). Before 2000, this alga was detected in Angara-Kichera Sor, Anga Bay, Barguzin Bay (settlement of Makarovo) and opposite the town of Baikalsk. In 2016, *Oedogonium* was also recorded outside the habitats mentioned above, i.e. in Listvennichny Bay, the settlement of Angasolka, Cape Kotelnikovsky, Zavorotnaya Bay and Senogda Bay.

The bottom cover percent with filamentous algae, predominantly with *Spirogyra*, varied between 0 and 100% in different regions of the coastal area at a depth of over 2 m. Velocities varied at maximal wave lengths of 14 m and 21 m and periodicity of 3 s and 5.1 s, respectively, and at wave heights of 1–1.2 m: *U* from 0.03 to 6.2 m s^−1^ and *V*_sh_ from 0.07 to 0.15 m s^−1^. The orbital velocity in the zone of wave profile transformation (at a depth of 3.5–3.7 m) was on average 0.32±0.03 m s^−1^, 1.12±0.06 m s^−1^ in the zone of wave breaking (depths of 1.5–2.5 m) and 3.76±0.06 m s^−1^ in the surf zone at the beginning of surge (depth of up to 1.5 m). The boundaries of zones (Fig. 2*a*) were mobile, and their location depended on gale force at Lake Baikal. At the same wave parameters, *K_m_* in these zones was higher than 1, attesting to the mobility of bottom sediments.

PCA analysis showed that the main percentage of variability (80%) of the whole database was provided by the first two principal components. According to the first principal component, with 56% of total variability of the database, environmental factors such as depth, height, length and periodicity of waves as well as width of the coastal zone affected the cover percent; the load of variables was positive (Fig. 4*a*). According to the second principal component, with 24% of total variability of the database, the distribution of filamentous algae was dependent on shear velocity, composition of bottom sediments, depth and slope angle of the bottom in the coastal zone (Fig. 4*b*). It is clear that the bottom cover percent with filamentous algae was independent of the water flow pressure on the vertical surface, as filament strands stretched along the bottom horizontally or fluctuated synchronously with the reciprocating motion of water flow. The load of variables (x_12_–x_18_ at different wave heights) on the first and second components were negative. The water flow pressure on the vertical surface at a wave height of 1–1.2 m was 11–72 g m^−2^ at a depth of 10–1.5 m and was insufficient for detachment of filaments from the substrate (underwater video filming).

**Fig. 4.**
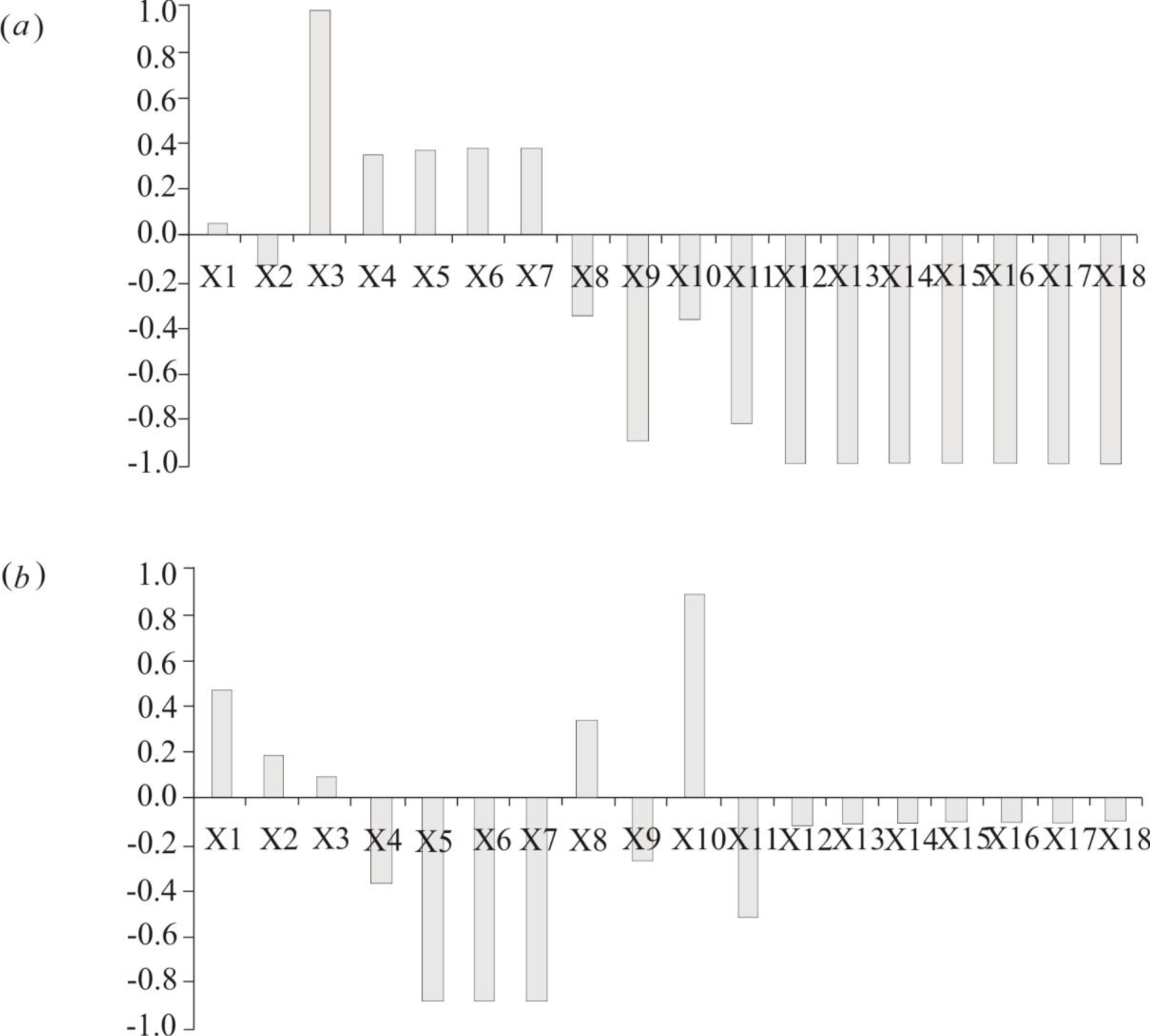
Loads of variables on the first (*a*) and second (*b*) principal components. Variables along X-axis: x_1_ – bottom cover percent with filamentous algae, %; x_2_ – composition of bottom sediments; x_3_ – depth, m; x_4_ – width of the coastal zone, m; x_5_ – wave height, m; x_6_ – wave length, m; x_7_ – periodicity of wave activity, c; x_8_ – slope ratio of the coastal zone; x_9_ – bottom current velocity, m∙s^−1^; x_10_ – shear velocity of sediment movement (0.5 mm in diameter), m∙s^−1^; x_11_ – coefficient of sediment mobility; x_12_-x_18_ – hydrodynamic pressure (g m^−2^) on vertical surface at certain depths (0.5 m; 1.5 m; 2 m; 3 m; 4 m; 5 m). Along Y-axis contribution of each variable x to total variability of characteristics.

In space of the two first principal components, the point set is divided into two non-overlapping subsets *I* and *II* (Fig. 5). Subset *I* covers the points of sampling at stations where filamentous algae are often recorded. Moreover, at some stations (Cape Listvennichny, Krestovka Valley, near the outlet of the Angara River, Baikalsk and Ushakovka River), the bottom cover percent with filamentous algae at a depth of over 2 m reached 40–100%. At these stations, the vertical zoning of the spatial distribution of meio- and macrophytes was disturbed because of mass development of filamentous algae. The same subset covers the stations (settlement of Kultuk, settlement of Maximikha and Ushakovka River) where we found free-lying clumps formed by filamentous algae on the sandy bottom. In addition, the first subset covered the sites where the bottom cover percent was up to 15%. However, the vertical zoning of algal distribution was not disturbed.

**Fig. 5.**
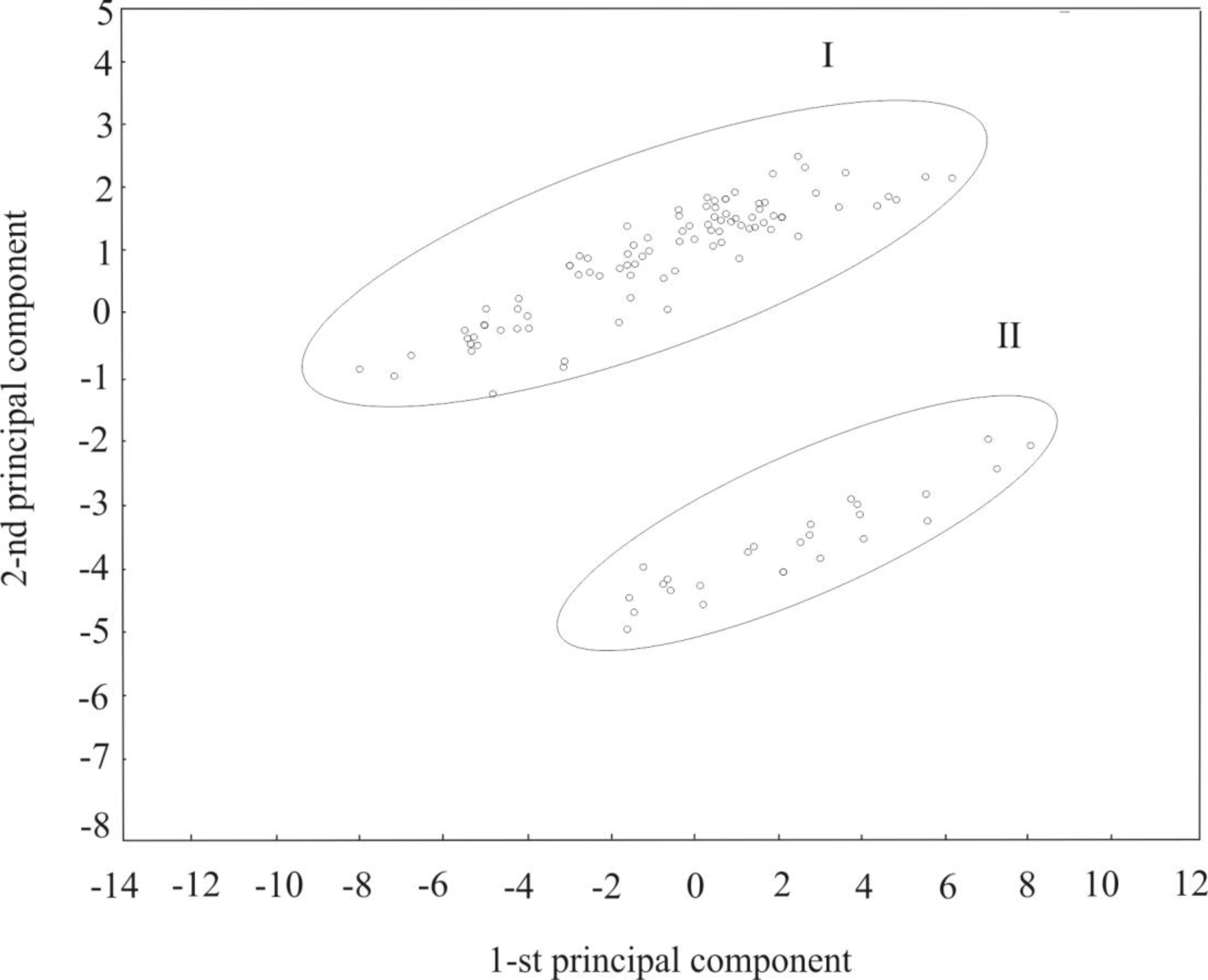
Location of sampling points in the space of two principal components. Subset *I* includes points where the bottom cover percent with filamentous algae was from 15 to 100%: Angasolka, Marituy, Cape Tolsty, near Angara River, Krestovka, Cape Listvennichny, Emelyanovka, Ushakovka, Khara-Murin and town of Baikalsk. Subset *II* covers points where there were no filaments or their cover percent was 1-3%: Cape Ryty, Cape Zavorotny, Cape Kotelnikovsky, Irinda Bay and Peninsula Svyatoy Nos.

Subset *II* comprises stations at which historically formed zoning in the spatial distribution of meio- and macrophytes remains; there were no filamentous algae or their cover percent was 1–3% (Fig. 5). The diversity and specific structure of meio- and macrophyte communities of the coastal zone were studied in the relatively uniform hydrodynamic environment at site 3 (Fig. 1*a*, Table 2) but with different recreation load within its boundaries. In 2016, in the background region opposite the Emelyanovka Valley, the Shannon index was not high (1.0). We revealed an algal community dominated by an endemic species *Draparnaldioides baicalensis* C. Meyer et Skabitsch. (Fig. 6*a*). Its composition was represented by 22 taxa with a total biomass of 112±58 g m^−2^. The filamentous alga *U. zonata* is characteristic of stony substrates in Lake Baikal. Singular filaments of three *Spirogyra* morphotypes were recorded among rare species (P<10%). Their biomass was low beyond the limits of balance sensitivity. The percentage of all filamentous algae was lower than 0.1% of the total biomass. In 1987, the Shannon index was 1.7. At this time, the community consisted of 15 taxa with the dominance of *D. baicalensis* (Fig. 6*b*). Members of the genus *Spirogyra* were absent among subdominant and minor species in the algal community. The total biomass of the community made up 90±27 g m^−2^, and only *U. zonata* was recorded in this community with a biomass of 2%.

In 2016, the Shannon index was 1.3 in the zone of anthropogenic impact opposite the settlement of Listvyanka (Krestovka Valley). The community with the dominance of *Spirogyra*, uncharacteristic of the open coastal zone of Lake Baikal, formed for the first time here (for the period of observations since the beginning of the previous year) (Fig. 6*a*). This community was represented by 25 taxa with a total biomass of 109±66 g m^−2^. Among minor species of filamentous algae in the community we registered *U. zonata* as well as members of the genus *Oedogonium* (two taxa), uncharacteristic of the open coastal zone in Lake Baikal. The contribution of filamentous algae in the total biomass of the community was 58%. Earlier (in 1987), the Shannon index (2.5) was twice as high. The community dominated by *Dermatochrysis reticulata* C. Meyer (Fig. 6*b*) included 23 taxa of meio- and macrophytes with a total biomass of 66±24 g m^−2^. The percentage of filamentous algae was 13% of the total biomass of the community, among which there were only members of the genus *Ulothrix* (*U. zonata, U. tenerrima* Kütz. and *U. tenuissima* Kütz.); *Spirogyra* and *Oedogonium* were not recorded.

**Fig. 6.**
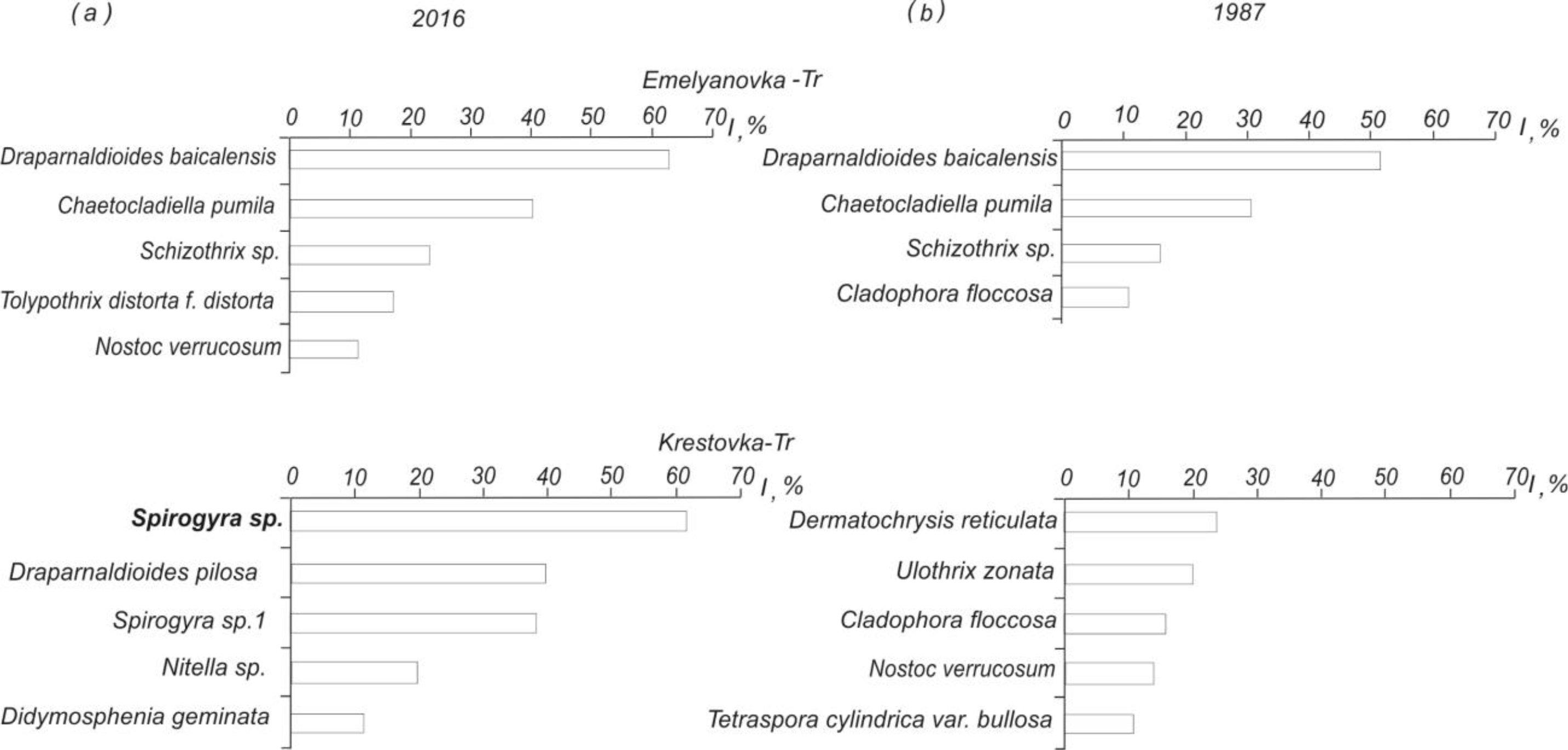
Structure of meio- and macrophytes communities in the coastal zone of Lake Baikal in the area of local anthropogenic impact (Krestovka Valley, Listvennichny Bay) and background area (Emelyanovka Valley) during different years (*a*) – in 2016; (*b*) – in 1987. Along X-axis – density index; along Y-axis – species ranking in the order of decrease of density index.

The distribution of *Spirogyra* biomass depends on the hydrodynamic environment. Its lowest content was recorded in the surf and wave breaking zones, whereas the highest values were registered in the zone of weak effect of wave activity on the bottom (Fig. 2*b*). Moreover, the bottom cover percent with filamentous algae increased in the bay with the depth increase, i.e. with the decrease of wave effect on the bottom (Fig. 2*c*).

According to the visual observations of I. Khanaev during the year, the length of *Spirogyra* thallomes varied in the coastal zone of Listvennichny Bay. In January, it was 7–10 cm and up to 20 cm in May. In June–August, filament strands could reach more than 150 cm and 10 cm in September–December. The longest filament strands usually form at depths below 8 m (up to 15 m), i.e. beyond the zone of wave effect on the bottom during summer storms (*h*=1 м). During our investigation, the length of filament strands reached 50–70 cm in some areas of the lake. Cells forming the *Spirogyra* thallomes were represented by two size groups. Some cells (n=38 measurements) were on average 30±1 μm wide (14–40 μm) and 152±14 μm long (29–345 μm). Others (n=23) were an average of 45±1 μm wide (41–68 μm) and 208±20 μm long (81–378 μm). The mass development of *Spirogyra* affects the structural organisation of both benthic and planktonic algal communities.

## Discussion

The average directional transfer of Baikal waters forms mainly under dynamic influence of the atmosphere (wind regime and pressure gradient above the water area). The system of currents at Lake Baikal (Fig. 1*b*) comprises alongshore circulation of cyclone type covering the entire lake and secondary circulations in the southern, central and northern basins of Lake Baikal (Afanasyev and Verbolov 1977). In the storm period, orbital velocities with a regime probability of 0.1% near the bottom can stir up the sand around the entire area of the coastal zone (Fialkov 1983). The system of currents and turbulent diffusion distribute terrigenous suspended sediments brought with river waters around the water area of the lake. According to a mathematical model (Mizandrontsev and Sudakov 1981), suspended sediments (diameter of 0.005 mm, density of 2.65 g cm^−3^ and sinking velocity of 17×10^−3^ mm∙s^−1^) are transported along the western coast of Northern Baikal at current velocities of some centimeters per second for a distance of hundreds of kilometers from the mouths of large tributaries located in the northern part of the lake. Offshore secondary circulations and horizontal turbulent diffusion promote the removal of suspended particles in the open areas of the lake (Mizandrontsev and Sudakov 1981). The diatoms *A. formosa* and *A. baicalensis* as well as fine mineral sediments can be transferred by coastal currents at significant distances. This relates to planktonic algae without exoskeletons with densities close to the density of the lake water and to fragments of *Spirogyra* thallomes. Moreover, the density of living skeletonless planktonic algae is close to 1 g cm^−3^ and can be lower than the water density (at this temperature) due to the presence of gas vacuoles and fat inclusions (Henderson-Sellers 1987; Smith 1982). The transfer mechanism of filaments and their fragments within the water column, considering the geometric and hydraulic characteristics (Table 1), is similar to that of planktonic diatoms. The deposition rate of *A. baicalensis* (10,000 μm^3^) in the laminar water flow is 39×10^−3^ mm s^−1^ (Votintsev 1961) and that of sea phytoplankton, particularly dinoflagellates (taking into account the equivalent radius of cells and their non-spherical shape), is from 3×10^−3^ mm s ^−1^ to 45×10^−3^ mm s^−1^ (Kamykowski *et al.* 1992). The deposition rate of *Spirogyra* (22×10^−3^ mm s^−1^) is lower than that of Baikal diatoms. As deposition rates are very low, it helps both filament fragments and diatoms remain within the water column for a long time. Thallome fragments of filamentous algae removed by currents from the coastal area can move along the perimeter in each basin of the lake and around the entire lake (Fig. 1*b*). For example, in the southern basin of Lake Baikal, algal fragments will be transferred from Cape Listvennichny to Cape Tolsty (Fig. 1*a*) for 11 days at the wind with regime probability of 50% and at the velocity of drift current of 9 cm s^−1^ in the middle water layer.

The growth and development of benthic algae in aquatic ecosystems are also closely connected with hydrodynamics, in particular with wave activity (Reiter 1986; Raspopov *et al*. 1990; Nozaki *et al*. 2003; Engelen *et al*. 2005). In Lake Baikal, as in other water bodies, vegetation of dominant algae of the vegetation belts (*U. zonata* at a depth of 0–1.5 m, *T. cylindrica* (Wahl.) Ag. var. *bullosa* C. Meyer and *Didymosphenia geminata* (Lingb.) M. Schmidt. at a depth of 1.5–2.5 m and species of the genus *Draparnaldioides* at a depth from 3 m to 10–12 m) is determined by the hydrodynamic environment. A large amount of floating fragments of algae and higher plants as well as their mass clumps on the shore of Lake Baikal after summer and autumn storms has been recorded since the beginning of the first half of the previous century (Kozhov 1931; Votintsev 1961). The detachment of benthic algae from the substrate can occur under the influence of orbital and reciprocating water motions caused by wave activity (Karabanov and Kulishenko 1990). In late autumn and early winter (November–December), when the waves reach their maximal height, the algae abovementioned stop their vegetation (Izhboldina 2007). It is clear that intense development of seasonal algae occurs in summer with durable calm and low wave activity (*h*=0.5 m). At this time, algal mats of filamentous algae can be found in the coastal zone at a depth of 3 m and deeper (Kravtsova *et al*. 2014).

Judging from in-situ data, in summer, *Spirogyra* forms the longest filament strands at depths of over 8–10 m. In Lake Baikal during rare summer storms with a wave height of about 1 m, bottom currents (*U_max_*=0.03–0.10 m s^−1^) are unable to detach filaments from the substrate, whereas at lower depths their detachment is quite probable. The filamentous algae *Spirogyra* and *Mougeotia* develop at current velocities of 0.12 m s^−1^ and 0.29 m s^−1^ with greater biomass in the first case and slightly lower biomass in the second case (Peterson and Stevenson 1992). In the majority of the studied regions of Lake Baikal, the orbital velocity (0.36 m s^−1^) emerging during the storm (*h*=1.0 m) at depths of 3.3–4 m is not critical for *Spirogyra* development in comparison with the areas where the velocity at these depths can reach 0.68 m s^−1^ (Fig. 5). Clumps of filaments freely lying on the sandy bottom after a 3-ball storm found during the field studies confirmed the wave effect on algae. In the Ushakovka River at a depth of 6 m (*U_max_*=0.18 m s^−1^), the filamentous clumps were likely formed from algae detached at 1.5–2.1 m, where current velocities could reach 1.02–3.80 m s^−1^. Depending on the slope angle and width of the coastal zone, algae detached from the substrate after the storm are either transferred by near-bottom currents or washed ashore or thrown to the shore. Freely lying *Spirogyra* clumps at the bottom of the coastal zone of Lake Baikal have been also found by other researchers (Timoshkin *et al*. 2016).

At lower depths (0.6–1.5 m), current velocities are significantly higher and can be 3.8–6.2 m s^−1^ (at *h*=1.0 m). At such velocities, coarse pebbles and boulders (30 cm in diameter) are transported and the length of algal filaments shorten, e.g. in *U. zonata* from 10–15 cm to 5–6 mm (Karabanov and Kulishenko 1990). Therefore, the main limiting factor in the development of filamentous algae in the zones of wave breaking and surge is mobility of bottom sediments. *Spirogyra* biomass here is several times lower than at a depth of 3 m (Fig. 2*b,c*) because of the abrasive effect of sandy particles (with eroding velocities minimal for binder soil) and gravel-pebble material coming into motion. In the coastal zone of seas and large lakes, orbital velocities (0.5 m s^−1^) are able to resuspend fine sand particles (Rasmussen and Rowan 1997). The movement of debris (1–20 cm in diameter) in water bodies is known to emerge when near-bottom current velocity exceeds critical values (*V_sh_*=0.5–1.7 m∙s^−1^) and depends on its size and bottom slope (Volkov and Ionin 1962). In Lake Baikal, as in other water bodies, bottom sediments are free from algae as a result of transfer, mobility, friction and turbidity of debris of different sizes. Filaments detached from the substrate enter the water column. This is supported by findings of *Spirogyra* in the planktonic samples at all studied depths in Southern Baikal (Fig. 3*a*).

Factors affecting spatial heterogeneity of benthic algae are numerous and one of them is anthropogenic impact. The intense development of filamentous algae in the coastal zone of Lake Baikal is caused by nutrient flux in the areas of human activity (Kravtsova *et al*. 2014; Timoshkin *et al*. 2016). In particular, in July–August 2011 high concentrations of nutrients P up to 0.420 mg L^−1^ and nitrate as N up to 0.20 mg L^−1^ were recorded in bottom waters of Listvennichny Bay, whereas the background values for P and N in open parts of the lake are 0.007 mg L^−1^ and 0.08 mg L^−1^, respectively (Khodzher *et al.* 2017). In addition, in 2015 the content of nutrients (mg L^−1^) was higher in comparison with that in the open lake: NH_4_^+^=0.560 ± 0.47; NO_2_^−^=0.055 ± 0.01; NO_3_^−^=0.690 ± 0.07; P _mineral_=0.025 ± 0.03; P _organic_=0.024 ± 0.02; P_total_=0.049 ± 0.05 (Kulakova *et al*. 2017). Higher concentrations of nutrients in the depression topography are caused by secondary pollution produced by algae degrading in the coastal zone opposite the settlement of Listvyanka.

Despite *Spirogyra* having been registered earlier in the coastal-sor zone (Fig. 1*c*), such quantities of this alga (Fig. 2*a*; Kobanova *et al*. 2016) have not been recorded in either benthic (Izhboldina 2007) or planktonic algal communities (Kozhov 1931; Popovskaya 1977). This means that the specific character of the recent structure of Baikal algal communities in comparison with that in the previous century is the presence of *Spirogyra* in their composition (Table 2, Fig. 3*a*, Fig. 6*a*). These algae have been found ubiquitously in the benthic communities of the studied habitats, except station 17 opposite Irinda Bay, an area characterised by elevated velocities of bottom currents. In addition, secondary circulations existing in Northern Baikal do not cover this area (Fig. 1*b*). Moreover, of interest are earlier findings of singular *Spirogyra* filaments outside the coastal zone at depths of 40– 80 m (Fig. 1*c*). Their emergence in the deep zone as well as the emergence of the thermophilic diatom *A. formosa* at depths of 300–600 m may be attributed to the run-off of warm waters along the slope at the boundary of the thermal bar (Likhoshway *et al*. 1996).

## Conclusions

Hydrodynamics plays an important role in the formation of the recent structure of algal communities of Lake Baikal. Under conditions of global warming, anthropogenic impact and flux of nutrients into the coastal zone, we observe the bottom overgrowing with filamentous algae *Spirogyra* and *Oedogonium*. During storms, filaments detach from the substrate and are washed ashore, forming aggregates on the beaches or entering the water column. For the hydraulic characteristics of filamentous algae, *Spirogyra* in particular are comparable with those of planktonic diatoms, and the existing system of currents in Lake Baikal causes their transfer from the regions of mass development and distribution outside the zones of local anthropogenic impact. Therefore, in the open parts of the coastal zone of Lake Baikal remote from settlements, we find only singular filaments or thallome fragments of *Spirogyra*. Bays and sors, i.e. traditional habitats of this alga, also serve as a source of replenishment with algal fragments of circular currents. Only at present has *Spirogyra* been registered in the plankton of the open parts of Lake Baikal despite benthic diatoms being constantly recorded in the composition of plankton. In the case of rising nutrient load on the coastal zone, the role of filamentous algae in the Baikal ecosystem will increase and hydrodynamics will promote its dissemination. The investigation of hydrodynamic processes in the interconnection with a biotic component of aquatic ecosystems plays an important role in the understanding of mechanisms of their function. The algal bloom in the inland waters has become a critically important issue for its impacts on natural and social environments. Long-term monitoring must therefore consider the human factor controlling these blooms and their impact on water supply in Lake Baikal and other large lakes threatened by accelerating eutrophication.

## Acknowledgements

This work was supported by budget projects of the Federal Agency of Scientific Organizations Nos. 0345-2016-0004 and 0345-2016-0006. Verification of meio- and macrophyte taxa with the use of molecular research techniques was carried out with the assistance of a grant Nos.17-44-388071 r_a. The authors thank divers V. Chernykh and Yu. Yushuk for their help in sampling.

